# Structural Characterization of mRNA Lipid Nanoparticles in the Presence of Intrinsic Drug-free Lipid Nanoparticles

**DOI:** 10.1101/2024.09.27.614859

**Authors:** Xiaoxia Chen, Yongfeng Ye, Mengrong Li, Taisen Zuo, Zhenhua Xie, Yubin Ke, He Cheng, Liang Hong, Zhuo Liu

**Affiliations:** School of Physics and Astronomy, Shanghai Jiao Tong University, Shanghai 200240, China; Institute of Natural Sciences, Shanghai Jiao Tong University, Shanghai 200240, China; Spallation Neutron Source Science Center, Dongguan 523803, China; Institute of High Energy Physics, Chinese Academy of Sciences, Beijing 100049, China; Shanghai National Centre for Applied Mathematics (SJTU Center), MOE-LSC, Shanghai Jiao Tong University, Shanghai 200240, China; Zhangjiang Institute for Advanced Study, Shanghai Jiao Tong University, Shanghai 201203, China; Shanghai Artificial Intelligence Laboratory, Shanghai 200232, China

## Abstract

Lipid nanoparticles (LNPs) have emerged as a versatile platform for mRNA delivery across a range of applications, including disease prevention, cancer immunotherapy, and gene editing. Structural models of mRNA-containing lipid nanoparticles (mRNA-LNPs) have also been proposed based on characterization of samples by using various advanced techniques. Among these, small angle neutron scattering (SANS) has proven essential for elucidating the lipid distribution within mRNA-LNPs, a factor crucial to both their preparation and efficacy. However, recent findings suggest that the mRNA-LNP samples prepared via commercial microfluidic techniques may contain a substantial fraction of drug-free LNPs, casting doubt on the validity of earlier structural models. In this study, we employed contrast variation SANS to characterize both drug-free LNPs and our mRNA-LNP sample, and quantified the proportion of drug-free LNPs present to be ∼30% in our mRNA-LNP sample using nano flow cytometry. By removing the contributions of drug-free LNPs from the SANS data of our mRNA-LNP sample, we were able to precisely characterize the structure of mRNA-LNPs. Consequently, we proposed structural models for both drug-free LNPs and mRNA-LNPs. Notably, our analysis revealed similar lipid distributions and shell thicknesses between the two particle types, while the solvent content in mRNA-LNPs was significantly higher, leading to a larger core size. This work not only offers a method for accurately characterizing the structure of mRNA-LNPs, but also establishes criteria for selecting appropriate analytical techniques based on the structural parameters of interest. Therefore, our findings hold significant implications for the mechanistic understanding and quality control of mRNA-based vaccines.

**Significance:** Precise structural determination of mRNA-containing lipid nanoparticles (mRNA-LNPs) is vital for mechanistic insights into their preparation, delivery, immunogenicity, and storage, which are critical to the development of mRNA-based vaccines. However, most previous studies overlooked the substantial presence of drug-free LNPs within these samples. Here, we identified that approximately 30% of the nanoparticles in our mRNA-LNP sample were drug-free. By integrating contrast variation small angle neutron scattering (SANS) data from both drug-free LNPs and mRNA-LNPs, we developed structural models for both particle types, and provided a guidance for characterization technique selection based on concerned structural features. Beyond mechanistic insight on structure, our approach offers a robust method for quality assessment and process monitoring in mRNA-based vaccine production.

## Introduction

Lipid nanoparticles (LNPs) have proven to be versatile nanocarriers for delivering nucleic acids in a range of medical applications, including cancer immunotherapy (1, 2), metabolic regulation (3, 4), and prophylactic interventions (5, 6). A typical LNP formulation comprises an cationic ionizable lipid, a helper lipid, cholesterol, and a PEGylated lipid, each contributing uniquely to the stability, structure, encapsulation efficiency, and *in vitro* and *in vivo* interactions (7, 8). Specifically, the internal structure and surface morphology of nucleic acid-loaded LNPs vary with lipid composition, leading to differences in stability and therapeutic efficacy (9-12). To optimize the delivery of nucleic acids via LNPs, extensive structural characterizations have been conducted on small interfering RNA (siRNA)-loaded LNPs (13-17), mRNA-loaded LNPs (mRNA-LNPs) (12, 13, 18-20), and plasmid DNA-loaded LNPs (21, 22) using advanced techniques such as cryogenic electron microscopy (cryo-EM) (14, 15, 18), nuclear magnetic resonance (NMR) (13, 14, 23), small angle X-ray scattering (SAXS) (12, 14, 22, 24), and small angle neutron scattering (SANS) (12, 21, 22, 25, 26), etc (16, 17, 19, 20). These analyses have informed the development of structural models (12-14, 17, 19, 21-23, 25), which are critical for understanding the mechanisms of LNP mediated nucleic acid delivery and guiding the rational design of more effective LNPs (10, 27-29).

The structural characterization of lipid nanoparticles (LNPs) is challenging due to the similar elemental composition of their four lipid components, making it difficult to differentiate and determine their distributions. Isotope contrast variation small angle neutron scattering (SANS), however, offers a solution by exploiting the differences in neutron scattering length density between hydrogen and deuterium (30). For instance, M. Y. Arteta and collaborators successfully used contrast variation SANS to elucidate lipid distributions, water content, and mRNA copy number within a single LNP (12, 25). Similarly, Z. Li et al. utilized SANS to investigate acidification induced structural changes in plasmid DNA-loaded LNPs (21). It is worth noting that previous SANS analyses assumed a homogeneous payload distribution within LNPs, disregarding the possible presence of drug-free LNPs (12, 21, 25, 26, 31). This assumption is reasonable for siRNA-loaded LNPs, where each LNP typically encapsulates hundreds of siRNA molecules (16, 17, 32). In contrast, for mRNA with ∼2,000 nucleotides loaded LNPs, S. Li et al. (20) and T. Sych et al. (33) reported that 40% to 80% of the LNPs were drug-free, containing no mRNA. The substantial presence of drug-free LNPs in the mRNA-LNP samples raises concerns about the reliability of structural data obtained from SANS and calls into question the structural models proposed in earlier studies (12, 25, 26, 31). This issue has begun to gain recognition within the field. Gilbert et al. addressed the presence of drug-free LNPs by fitting SANS data of nucleic acid-loaded LNPs using a combined model of a core-shell sphere, supplemented by a broad peak representing the internal structure around 1 nm^-1^ (22). However, this model assumed that the lipid distributions and water content of drug-free and mRNA-loaded LNPs were identical, contradicting the structural models proposed by T. Unruh et al. (24), who suggested that these two nanoparticles have distinct water content. Importantly, even the structural information provided by T. Unruh et al. remains an average of drug-free and mRNA-loaded LNPs. Thus, the structural models derived from previous SANS studies warrant further re-evaluation in light of the significant proportion of drug-free LNPs in mRNA-LNP samples.

Herein, we encapsulated mRNA molecules containing 2,856 nucleotides into lipid nanoparticles (LNPs) and determined that approximately 30% of the LNPs in our mRNA-LNP sample were drug-free using nano flow cytometry (NanoFCM) (32, 34). We then employed contrast variation very small angle neutron scattering (VSANS) to characterize both drug-free LNPs and our mRNA-LNP sample. A structural model of drug-free LNPs was constructed based on the measurement of the drug-free LNP sample. More importantly, we developed a method to isolate the structural information of mRNA-LNPs by subtracting the VSANS data of drug-free LNPs from the mRNA-LNP sample, accounting for the proportion of drug-free LNPs. Through fitting the VSANS data, we determined key structural parameters of mRNA-LNPs, including lipid distribution, solvent content, and mRNA copy number per LNP, and constructed the corresponding structural model. As a result, we accurately quantified the proportions of drug-free LNPs and mRNA-LNPs within the sample, and established distinct structural models for both particle types. Our findings offer a refined structural characterization of mRNA-LNPs, taking into account for the intrinsic presence of drug-free LNPs. This provides a more accurate model for mRNA-LNP samples, which has important implications for the mechanistic study and quality assessment of nucleic acid-loaded LNPs.

## Results

### Basic characterizations of the drug-free LNP and mRNA-LNP samples

In this study, we utilized a lipid formulation developed by Moderna, consisting of the ionizable cationic lipid SM-102, cholesterol, distearoylphosphatidylcholine (DSPC), and PEGylated lipid (DMG-PEG2000) in a molar ratio of 50:38.5:10:1.5 (5). This formulation, used in the COVID-19 mRNA vaccine, is well known for its efficiency in nucleic acid encapsulation and intracellular delivery. mRNA molecules containing 2,856 nucleotides were encapsulated within this lipid formulation using a microfluidic technique (details in Materials and Methods). For comparison, a drug-free LNP sample was prepared using the same lipid formulation but without nucleic acids. As shown in Figure 1a, the mRNA-LNP sample exhibits a larger particle size compared to the drug-free LNP sample, although both display similar size distributions. Cryogenic electron microscopy (cryo-EM) images (Figure 1b) reveal that both drug-free LNPs and the mRNA-LNP sample exhibit a bleb-like morphology, i.e., an approximately spherical but irregular structure with distinct electron density partitions. This bleb morphology of mRNA-LNPs has been reported in several previous studies (18, 31, 34-36), despite most structural studies on LNPs have focused on spherical particles (12, 14, 25, 37). Based on the electron density differences observed in the bleb particles, we propose that our mRNA-LNPs are composed of a water compartment and a lipid compartment containing mRNA, consistent with the structural model proposed by Leung et al. (35).

**Figure 1.**
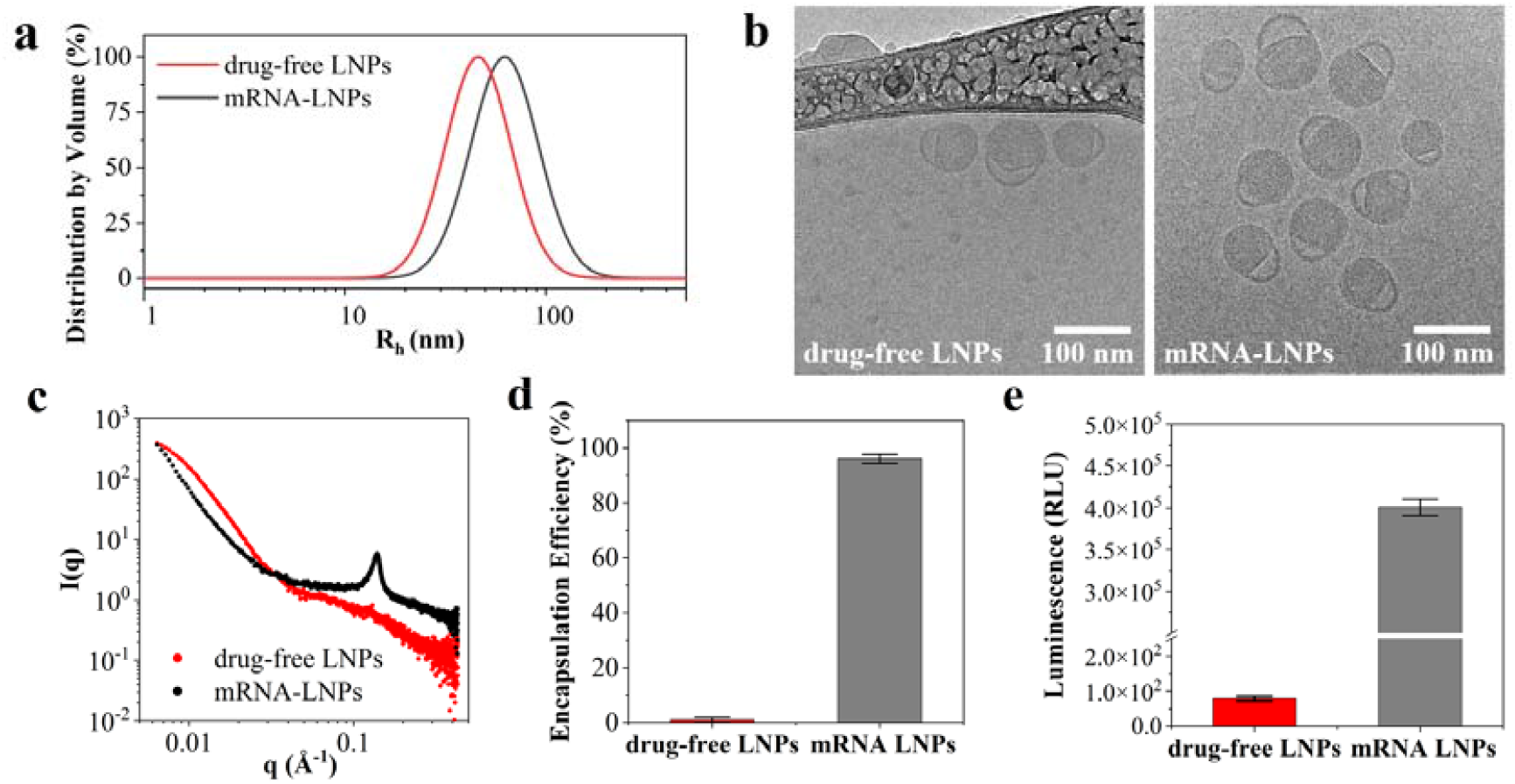
Basic characterizations of the drug-free LNP and mRNA-LNP samples. (a) Particle size distributions, (b) cryo-EM images, (c) SAXS curves, (d) encapsulation efficiency, and (e) *in vitro* transfection potency of the drug-free LNP and mRNA-LNP samples. HEK293T cells were transfected by the LNP and mRNA-LNP samples. Luciferase expression was quantified after 24 h of incubation (n = 3).

To further investigate the internal structure of the mRNA-LNPs, we characterized both the drug-free LNP and mRNA-LNP samples using small angle X-ray scattering (SAXS). The SAXS curve for the mRNA-LNP sample (Figure 1c) shows a peak at *q* ∼0.13 Å^-1^, corresponding to a correlation distance of approximately 5 nm, which is absent in the drug-free LNP sample. This result aligns with previous studies (12), where a peak around *q* ∼0.1 Å^-1^ was attributed to the characteristic distance between mRNA molecules embedded in a lipid matrix. Additionally, no peak was observed in the SAXS data for an mRNA aqueous solution (31), supporting the presence of a lipid-mRNA mixture in our mRNA-LNPs. This structural model differs from those in other studies, where the bleb morphology was attributed to mRNA-containing water compartments and lipid compartments (18, 31, 36). The radius of gyration (R_g_) of both samples was calculated from the SAXS data (Figure 1c) (38). As shown in Table 1, the R_g_ values for drug-free LNPs and mRNA-LNPs are approximately 32.8 nm and 46.1 nm, respectively, while their hydrodynamic radii (R_h_) are 47.8±2.0 nm and 63.5±0.6 nm. We further assessed the encapsulation efficiency and *in vitro* transfection potency of both samples (Figures 1d and 1e), confirming the high quality and efficacy of our mRNA-LNP formulation.

**Table 1.**
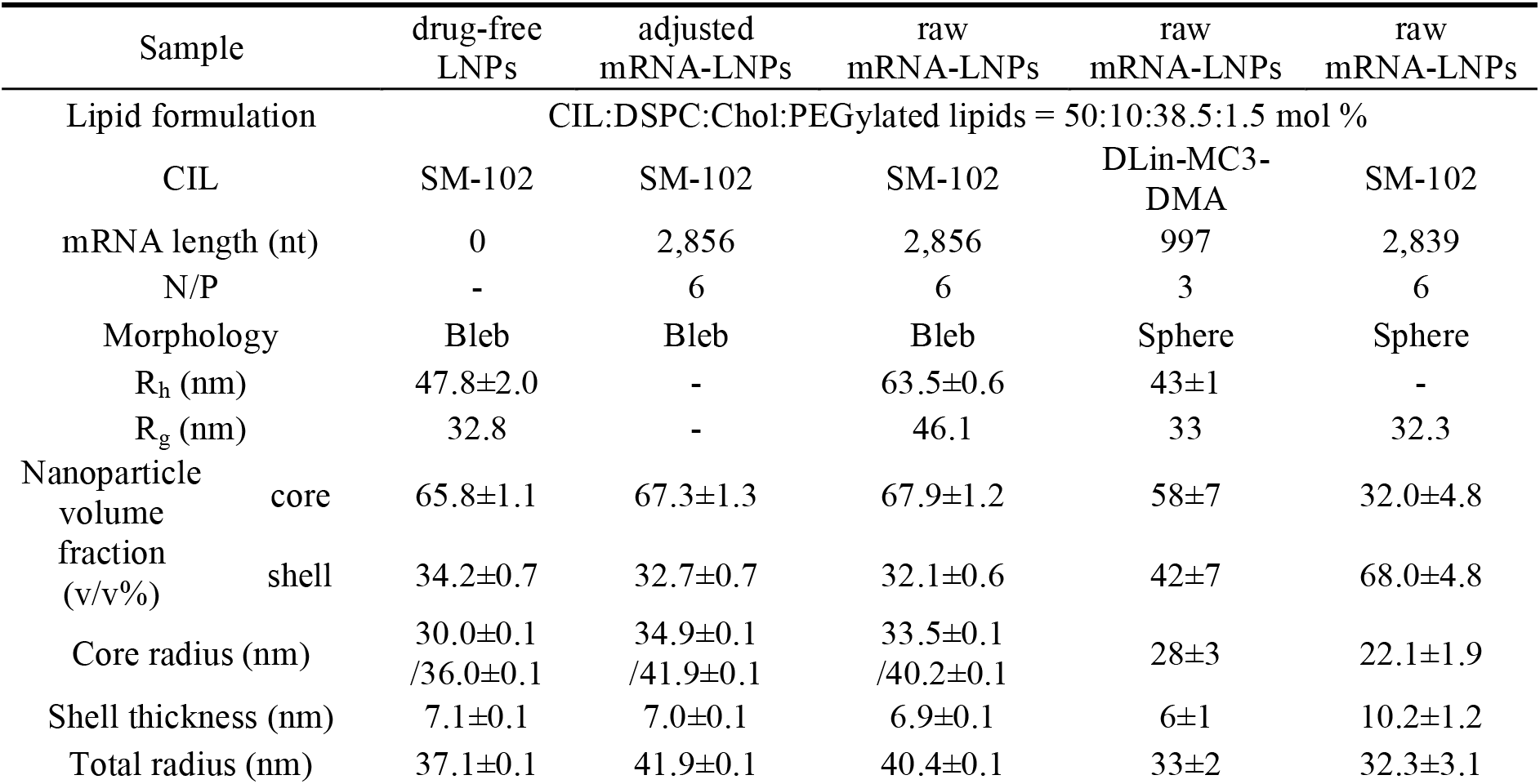
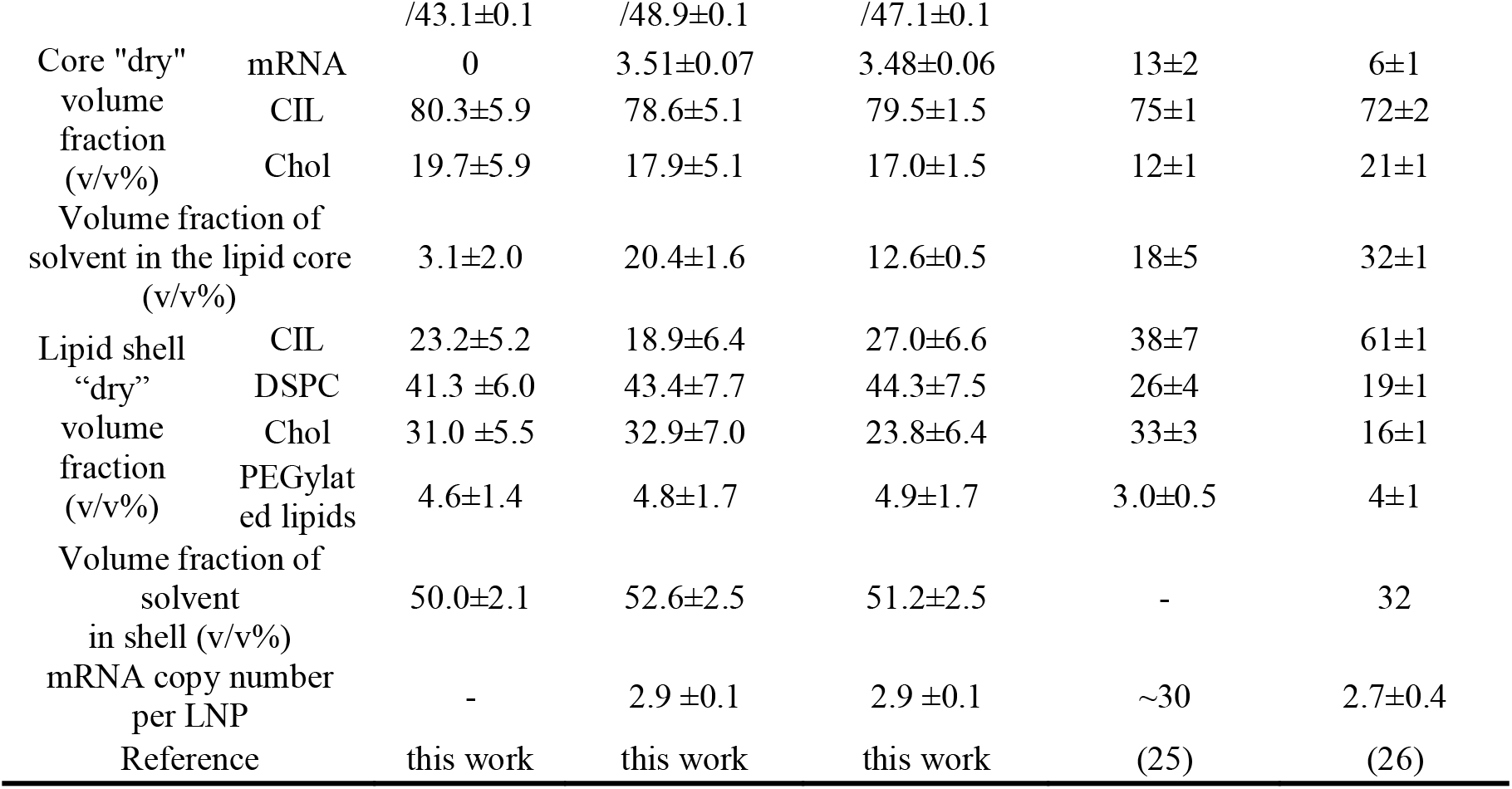
Structure information of drug-free LNPs and mRNA-LNPs determined by VSANS in this work, and structural parameters of the mRNA-LNP samples in Refs. (25) and (26).

Raw mRNA-LNPs refer to the experimental mRNA-LNP samples, comprising both drug-free LNPs and mRNA-loaded LNPs. In contrast, adjusted mRNA-LNPs represent mRNA-LNPs isolated from these samples by excluding the contribution of drug-free LNPs, containing only mRNA-loaded LNPs. The volume fractions were estimated from the fitted scattering length density (SLD) using a core-shell ellipsoid model. Abbreviations used include DLin-MC3-DMA (*O*-(*Z*,*Z*,*Z*,*Z*-heptatriaconta-6,9,26,29-tetraen-19-yl)-4-(*N*,*N*-dimethylamino)-butanoate), CIL (cationic ionizable lipid), DSPC (1,2-distearoyl-sn-glycero-3-phosphocholine), and Chol (cholesterol). The N/P ratio refers to the molar ratio of amine groups in the CIL to phosphate groups in mRNA. Hydrodynamic radius (R_h_) was determined by dynamic light scattering (DLS), and radius of gyration (R_g_) was measured via small angle X-ray scattering (SAXS) or small angle neutron scattering (SANS). In the structural analysis of mRNA-LNPs, the distributions of DSPC and DMG-PEG lipids were fixed within the shell, while mRNA was localized in the core. Given the bleb morphology of both drug-free and mRNA-loaded LNPs, the core and total radii of our samples are expressed as an equatorial radius followed by a polar radius, reflecting their ellipsoidal structure. Due to the presence of water compartment in our samples, the volume fraction of solvent in the lipid core was extracted to compare with previous results in Refs. (25) and (26). The error reported is from uncertainty analysis using the algorithm DREAM (38) and error propagation.

### Structural characterization of drug-free LNPs by VSANS

Lipid nanoparticles (LNPs) play dual roles in mRNA vaccines, serving as adjuvants to enhance efficacy (39) and as potential immunogens that may induce adverse effects (40). Therefore, understanding the structure of LNPs, particularly in cases where drug-free LNPs are present in significant amounts (20, 33), is critical for elucidating the pharmacokinetics and biodistribution of mRNA vaccines. Despite their importance, detailed structural characterizations of drug-free LNPs remain scarce. In this work, we applied contrast variation very small angle neutron scattering (VSANS) to characterize the structure of drug-free LNPs. The LNP samples contained partially deuterated d70-DSPC and d7-cholesterol, and were suspended in solvents with four distinct D_2_O/H_2_O ratios (30%, 51%, 67%, and 100% D_2_O) (Figure 2a). The VSANS data were collected in a *q* range from 0.003 Å^-1^ to 0.2 Å^-1^, enabling the investigation of lipid distributions within both the core and shell of the LNPs. By varying the degree of isotope contrast, we were able to selectively highlight different components of LNPs, facilitating a more precise determination of the structural arrangement and lipid distribution within the core and shell. This approach provides critical insight into the structural organization of drug-free LNPs, a previously unexplored area in the context of nucleic acid-loaded nanoparticle formulations.

**Figure 2.**
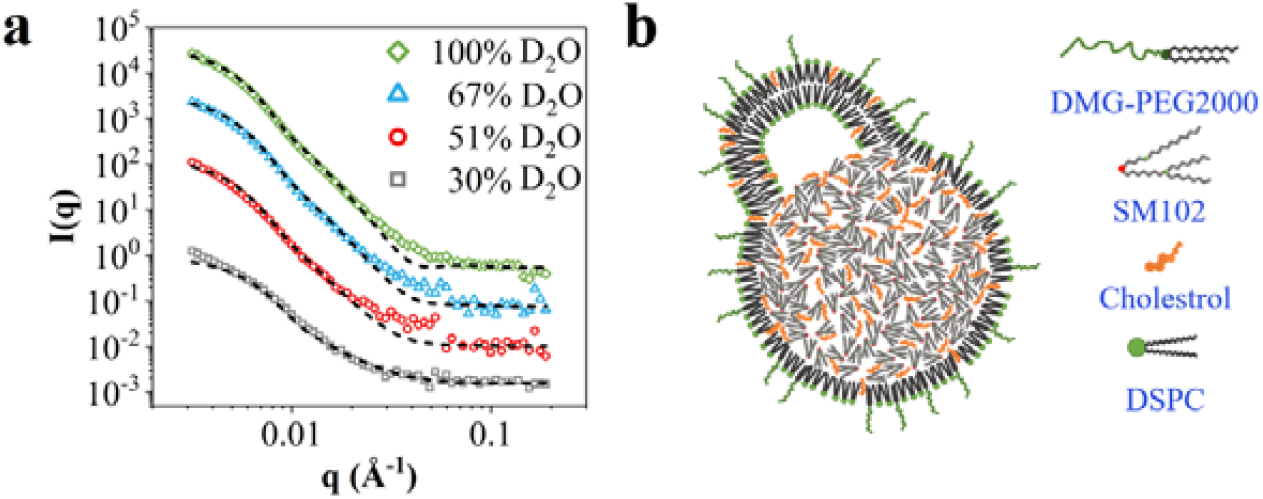
Structural characterization of drug-free LNPs by VSANS. (a) VSANS data collected for drug-free LNPs in four different solvent contrasts: 30% D_2_O (grey squares), 51% D_2_O (red dots), 67% D_2_O (blue triangles), and 100% D_2_O (green diamonds). The black dash lines are the results of model fitting. (b) The structural model of drug-free LNPs proposed according to the VSANS fitting results.

Figure 1b illustrates the morphological similarity between drug-free LNPs and mRNA-LNPs, both exhibiting a core-shell bleb structure. To model these structures, we applied a core-shell ellipsoid model (41) in SasView (https://www.sasview.org) to fit the four VSANS curves shown in Figure 2a. This model incorporates five fitting parameters: the equatorial radius of the core (r_equat_), the polar radius of the core (r_polar_), the shell thickness (t_shell_), the scattering length density of the core (SLD_core_), and the scattering length density of the shell (SLD_shell_) (see Figure S1 in Supporting Information, SI). Given that VSANS data were collected in solvents with only four distinct D_2_O/H_2_O ratios, we estimated the ratio between r_equat_ and r_polar_ based on cryo-EM images of drug-free LNPs and mRNA-LNPs (Figure 1b), fixing the ratio at 1.2 for our fitting. For each sample, the core radii and shell thickness were constrained to be consistent across the four solvent contrasts, while the SLD values of the core and shell were allowed to vary independently for each contrast. It is noteworthy that the core-shell ellipsoid model assumes homogeneity in the core’s SLD, which contrasts with the heterogeneous core structure observed in Figure 1b. To address this, we calculated the SLD of the mixed mRNA and lipids in the core using the following relationship: SLD_core_=(vf_core_-vf_mixed_) × SLD_sol_ + V_mixed_ × SLD_mixed_, where vf_core_ is the volume of the core, vf_mixed_ is the spherical volume of the mixed mRNA and lipids in the core, and SLD_sol_ and SLD_mixed_ are the scattering length densities of the solvent and the mixed components, respectively. In addition, we calculated the volume fractions of core and shell in the drug-free LNPs, along with the solvent content and lipid composition in both regions (Table 1). Further details regarding the VSANS data analysis are provided in the Materials and Methods section.

According to the structural information obtained from the VSANS data, we proposed a structural model for drug-free LNPs (Figure 2b). To validate our findings, we conducted additional SANS measurements using a different spectrometer in a *q* range of 0.005 Å^-1^ to 0.6 Å^-1^ at the China Spallation Neutron Source. As shown in Figure S2, the data collected from both VSANS and SANS exhibit similar behavior within the overlapping *q* range. However, due to limited beam time, the SANS experiments were performed only in solvents with D_2_O volume fractions of 30%, 51%, and 67%. Despite this limitation, the consistency between the two datasets supports the robustness of our structural model.

### The proportion of drug-free LNPs in the mRNA-LNP sample

To obtain the structural information of mRNA-LNPs, we need to figure out the proportion of drug-free LNPs in the mRNA-LNP sample, thereby we utilized nano flow cytometry (NanoFCM) to detect fluorescence and side scattering at the single nanoparticle level. Since the mRNA molecules were labeled with a fluorescent dye, both fluorescence and side scattering were observed for mRNA-LNPs, while only side scattering was detected for drug-free LNPs (Figure 3a). This allowed us to quantify the proportion of drug-free LNPs in the mRNA-LNP sample, which was found to be 27.6±2.9%, consistent with previous studies using the same technique (34). Additionally, we calculated the number-average payload of our mRNA-LNPs to be 2.9±0.1 mRNA molecules, each 2,856 nucleotides in length (Figure 3b). It is important to note that the number of nucleic acid molecules encapsulated in each LNP is highly dependent on their length. For shorter siRNA-containing LNPs, hundreds of siRNA molecules can be encapsulated within each LNP (16, 17, 32). For mRNAs with lengths of 1,029 and 2,839 nucleotides, the payloads were 2.80±0.41 (20) and 2.7±0.4 (26) per LNP, respectively. Therefore, it is expected that fewer than three mRNA molecules are encapsulated per LNP when the mRNA length exceeds 2,000 nucleotides.

**Figure 3.**
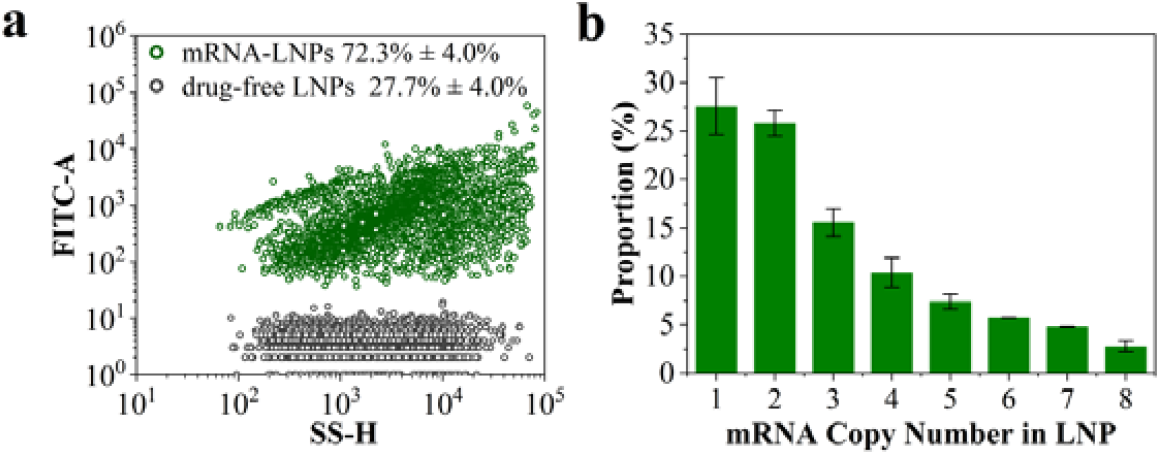
Determination of the proportion of drug-free LNPs in the mRNA-LNP sample. (a) Compiled bivariate dot-plot of the FL (fluorescence) burst area versus the SS (side scattering) burst area for drug-free LNPs (black circles) and mRNA-LNPs (green circles). (b) The mRNA copy number distribution in the mRNA-loaded LNPs.

### Structure of mRNA-LNPs characterized by VSANS

To elucidate the structure of mRNA-LNPs in the presence of intrinsic drug-free LNPs, we collected VSANS data for the mRNA-LNP sample containing deuterated DSPC (d70-DSPC) and deuterated cholesterol (d7-cholesterol) in solvents with varying D_2_O/H_2_O ratios. Figure 4 displays the VSANS profiles for the mRNA-LNP sample, composed of both mRNA-LNPs and drug-free LNPs (red circles and lines). Building on prior analyses, we had already determined the VSANS data for drug-free LNPs (Figure 2a) and estimated their proportion in the mRNA-LNP sample (∼30%). Using this information, we were able to subtract the contribution of drug-free LNPs from the VSANS data of the mixed mRNA-LNP sample, thereby isolating the structural signal of mRNA-LNPs. For clarity, we denote the mRNA-LNP sample with mixed drug-free LNPs and mRNA-LNPs as raw mRNA-LNPs, while mRNA-loaded LNPs in the mixed mRNA-LNP sample are denoted as adjusted mRNA-LNPs.

**Figure 4.**
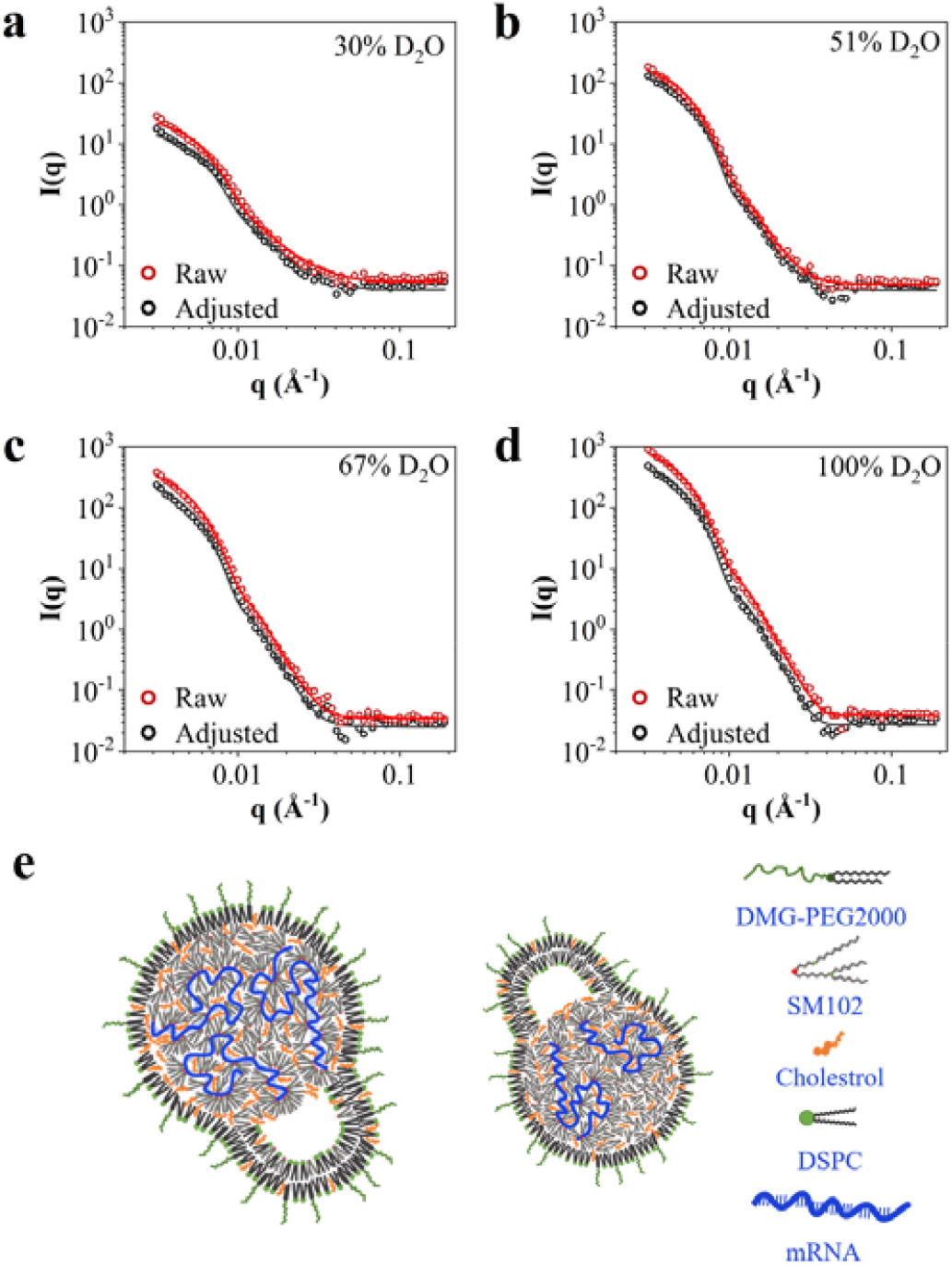
Structural characterization of mRNA-LNPs by VSANS. VSANS data collected for adjusted mRNA-LNPs (black circles) and raw mRNA-LNPs (red circles) in four different solvent contrasts: (a) 30%, (b) 51%, (c) 67%, and (d) 100% D_2_O buffer. Note that our raw mRNA-LNPs contained 70% mRNA-LNPs and 30% drug-free LNPs, and adjusted mRNA-LNPs correspond to the 70% mRNA-LNPs in the raw mRNA-LNPs. The black and red solid lines are the results of model fitting for the mRNA-LNP sample and mRNA-LNPs. (e) The structural model of mRNA-LNPs proposed according to the VSANS fitting results.

As shown in Figure 4, the VSANS data for adjusted mRNA-LNPs exhibit a distinct profile from raw mRNA-LNPs across all solvent contrasts. Similar to our approach with drug-free LNPs, we fit the VSANS data of mRNA-LNPs using a core-shell ellipsoid model. For consistency, we maintained the ratio of equatorial to polar radii at 1.2 and kept the radius and shell thickness constant across the different solvent contrasts. However, the scattering length densities (SLDs) of the core and shell were allowed to vary independently for each contrast. From the data fitting, we estimated the volume fractions of core and shell, as well as the solvent content and lipid composition within core and shell of mRNA-LNPs, and compared these values to those of drug-free LNPs (Table 1). Our analysis also indicated that the average number of mRNA molecules per LNP was 3.2± 0.2, in accordance with the payload determined by NanoFCM (Figure 3b). A detailed description of the VSANS data analysis is provided in Materials and Methods. Based on the structural parameters obtained from the fitting, we propose a refined structural model for mRNA-LNPs (see Table 1 and Figure 4e).

## Discussion

While the structure of mRNA-LNPs has been extensively investigated using small-angle neutron scattering (SANS), the presence of drug-free LNPs within the mRNA-LNP samples has often been overlooked, potentially introducing bias into existing structural models. This oversight is particularly significant given that the proportion of drug-free LNPs in samples prepared via commercial microfluidic techniques can exceed 40%. These drug-free LNPs may also play an important role in modulating immune responses, acting both as adjuvants and immunogens (42, 43). Thus, several critical questions arise: What is the structural model of drug-free LNPs? How can we accurately resolve the structure of mRNA-LNPs within a mixture that includes both drug-free and mRNA-loaded LNPs? Are the structures of drug-free LNPs and mRNA-LNPs comparable, or do they differ substantially? Finally, can we reliably infer the structure of mRNA-LNPs by examining the mixture of both types of particles, or does the presence of drug-free LNPs necessitate separate characterization to avoid structural misinterpretation? Addressing these questions is essential for advancing the structural understanding and biological assessment of mRNA-LNP systems.

To address these questions, we conducted contrast variation very small-angle neutron scattering (VSANS) on both drug-free LNPs and our mRNA-LNP sample. The mRNA-LNP sample is the mixture of both drug-free and mRNA-loaded LNPs, denoted as raw mRNA-LNPs in this study. In parallel, we quantified the proportion of drug-free LNPs in the raw mRNA-LNPs using nano flow cytometry (NanoFCM). By subtracting the contribution of drug-free LNPs from the VSANS data of raw mRNA-LNPs, we were able to isolate the structural data specific to mRNA-LNPs. Here we denoted mRNA-LNPs as adjusted mRNA-LNPs to distinguish them with raw mRNA-LNPs. Structural parameters for both drug-free LNPs and adjusted mRNA-LNPs were then extracted using a core-shell ellipsoid model (Table 1), with the corresponding structural models shown in Figures 2b and 4e, respectively. Also, we developed a methodology to resolve the structure of mRNA-LNPs within a mixture of drug-free and mRNA-loaded LNPs. This approach holds significant relevance for mechanistic studies related to the formulation (22, 26, 44), delivery (12, 25, 45, 46), immunogenicity (47, 48), and storage (49, 50) of mRNA-based vaccines, offering a more precise understanding of their structural dynamics and potential functional implications.

As shown in Table 1, the shell thickness and lipid distributions of drug-free LNPs and adjusted mRNA-LNPs are notably similar. The shell thickness (∼7.1 nm) approximates the bilayer thickness of DSPC (5.8 nm) (51), suggesting that a disordered bilayer may form due to the high curvature induced by SM-102. Although Arteta et al. demonstrated that lipid distribution in the shell of mRNA-LNPs can vary with different lipid compositions (12), our results indicate that the shell structure in LNPs remains independent of mRNA encapsulation. Moreover, the lipid distribution within the core is comparable between drug-free LNPs and adjusted mRNA-LNPs, though the core size is larger in adjusted mRNA-LNPs. Given that mRNA occupies only ∼4% of the core volume in adjusted mRNA-LNPs, we attribute the larger core size to the increased water content introduced by the hydrophilic mRNA molecules. The solvent volume fractions in the lipid cores of drug-free LNPs and adjusted mRNA-LNPs are ∼3% and ∼20%, respectively. As a consequence, we suggest that water in both drug-free LNPs and adjusted mRNA-LNPs remains non-freezing at low temperature (52, 53). However, the water content of ∼20% in the lipid core of mRNA-LNPs could induce osmotic stress during freezing, potentially causing particle rupture and reducing efficacy (49). The additional water in adjusted mRNA-LNPs also increases the likelihood of hydrolysis of both mRNA and cationic ionizable lipids (54-56). Lyophilization, which removes water molecules, may thus offer a promising strategy for the long-term storage of mRNA-LNPs (50, 57-59). Overall, the lipid distributions between drug-free LNPs and adjusted mRNA-LNPs are largely consistent, but adjusted mRNA-LNPs encapsulate additional mRNA and water molecules compared to drug-free LNPs.

Furthermore, we sought to determine whether the structural characteristics of mRNA-LNPs can be approximated by characterizing raw mRNA-LNPs. The contrast variation VSANS data for raw mRNA-LNPs were fit using the core-shell ellipsoid model, as depicted in Figure 4. Note that most structural parameters between adjusted and raw mRNA-LNPs were comparable, with the primary difference observed in the volume fraction of solvent within the lipid core (Table 1). Although a slightly smaller lipid core size and reduced mRNA volume fraction were observed in the lipid core of raw mRNA-LNPs, these differences were not statistically significant. Given the similarity in lipid distributions between drug-free LNPs and adjusted mRNA-LNPs, we anticipate that the lipid distribution in raw mRNA-LNPs closely mirrors that of mRNA-LNPs. Notably, the volume fraction of solvent in the lipid core of adjusted mRNA-LNPs was 20.4%. In contrast, the volume fraction was only 12.6% when fitting the VSANS data for raw mRNA-LNPs, suggesting that the solvent content in the lipid core of mRNA-LNPs could be substantially underestimated if the presence of drug-free LNPs is ignored. Moreover, we suspect that both lipid core size and mRNA copy number per LNP may be underestimated when the proportion of drug-free LNPs exceeds 30%. However, the shell thickness and lipid distributions in both core and shell appear independent of mRNA encapsulation, indicating that they can be reliably estimated by characterizing the mixture of drug-free LNPs and mRNA-LNPs. These insights allow us to tailor the characterization approach based on the structural parameters of interest. For example, shell thickness and lipid distribution in mRNA-LNPs can be obtained through contrast variation SANS on the mixture of drug-free LNPs and mRNA-LNPs. On the other hand, if lipid core size and mRNA payload per LNP are the focus, the proportion of drug-free LNPs in the sample should ideally be less than 30%. Importantly, determining the solvent content in mRNA-LNPs is highly sensitive to the amount of drug-free LNPs, underscoring the need for simultaneous SANS measurements on both raw mRNA-LNPs and the proportion of drug-free LNPs within the mixed sample.

## Conclusion

In summary, we have developed a method for characterizing the structure of mRNA-LNPs in the presence of drug-free LNPs by integrating contrast variation SANS and NanoFCM. From the SANS data, we generated a structural model of drug-free LNPs, providing valuable insights into their potential role in immune responses. Using NanoFCM, we quantified the proportion of drug-free LNPs in our mRNA-LNP sample, which was approximately 30%. This allowed us to isolate the SANS curves specific to mRNA-LNPs by subtracting the contributions of drug-free LNPs from the composite SANS data. Moreover, we resolved the structural features of mRNA-LNPs within a mixture of drug-free LNPs and mRNA-LNPs, proposing an accurate model for mRNA-LNPs. It is well-established that the lipid distribution in mRNA-LNPs plays a critical role in their *in vitro* and *in vivo* efficacy (10, 12, 25, 27) and their structural evolution during preparation (22, 26, 44). Furthermore, variations in the internal structure of mRNA-LNPs can influence the formation of a protein corona, which in turn affects delivery efficiency (25, 45, 60). Therefore, precise structural characterization of both drug-free LNPs and mRNA-LNPs is essential for the rational design and optimization of mRNA vaccines.

Additionally, we realized that the lipid distributions and thickness of shell were quite similar by comparing the structural parameters between drug-free LNPs and mRNA-LNPs. Therefore, the two structural information can be acquired via SANS, regardless of the amount of drug-free LNPs in the sample. In contrary, the solvent content in drug-free LNPs is much less than that in mRNA-LNPs. The higher solvent content in mRNA-LNPs also give rise to a larger lipid core size than that of drug-free LNPs. However, we note that the lipid core size and mRNA payload per LNP can be approximated by directly characterizing the mRNA-LNP sample only if the proportion of drug-free LNPs is less than 30%. Our methodology not only enables the determination of mRNA-LNP structure in the presence of drug-free LNPs, but also establishes a framework for selecting appropriate characterization techniques based on the specific structural parameters of interest.

## Acknowledgements

This work was supported by the National Natural Science Foundation of China (12204302), the Shanghai Pujiang Program (Grant No. 22PJ1406900), the Startup Fund for Young Faculty at SJTU (SFYF at SJTU), the Oceanic Interdisciplinary Program of Shanghai Jiao Tong University (Project No. SL2022MS018), the Natural Science Foundation of Shanghai (Grant No. 23ZR1431700), and the Student Innovation Center at Shanghai Jiao Tong University. We would like to thank Dr. Na Li from BL19U2 beamline of Shanghai Synchrotron Radiation Facility (SSRF) for the help with synchrotron small angle X-ray scattering measurements on No. 2022-NFPS-PT-007003.

## Author contributions

Z.L. and L.H. designed and supervised the project. X.C. and M.L. prepared the samples for neutron scattering and SAXS, and performed the measurements. Z.X. and Y.K. contributed to the small-angle neutron scattering experiments at BL-01 at China Spallation Neutron Source (CSNS) in China. T.Z. and H.C. assisted the very small angle neutron scattering at BL-14 at CSNS in China. X.C., Y.Y., and Z. L. did the data analysis. X.C., Y.Y., L.H., and Z.L. wrote the manuscript.

## Additional information

Correspondence and requests for materials should be addressed to.

## Materials and Methods

### Materials

An enhanced green fluorescent protein-linked firefly luciferase (eGFP-fLuc) encoding mRNA with sequence length of 2,856 nucleotides, was obtained from Yaohaibio Co., Ltd. (Jiangsu, China). This mRNA was synthesized via *in vitro* transcription, incorporating a Cap1 structure at the 5’ end, pseudouridine substitutions, and a 150-nucleotide poly(A) tail, with a purity of greater than 95% for the full-length construct. The cationic ionizable lipid (CIL), heptadecan-9-yl 8-((2-hydroxyethyl)(6-oxo-6-(undecyloxy)hexyl)amino)octanoate (SM102, CAS: 2089251-47-6), was purchased from Sinopeg Biotechnology Co., Ltd. (Fujian, China). 1,2-Distearoyl-sn-glycero-3-phosphocholine (DSPC, CAS: 816-94-4) was sourced from Merck KGaA (Darmstadt, Germany). Cholesterol (Chol, CAS: 57-88-5) and 1,2-Dilauryl-rac-glycero-3-methoxypolyethylene glycol 2000 (DMG-PEG2000, CAS: 160743-62-4) were obtained from Avanti Polar Lipids, Inc. (Alabaster, AL, USA). Trehalose (CAS: 6138-23-4) was purchased from Shaoxin Biotechnology Co., Ltd. (Shanghai, China).

### Sample Preparation

mRNA was dissolved in a 100 mM citrate buffer (pH 4.0) at a concentration of 72 μg/mL, serving as the aqueous phase. The lipid mixture, composed of SM-102, cholesterol, DSPC, and DMG-PEG2000 in a molar ratio of 50:38.5:10:1.5, was dissolved in anhydrous ethanol. Both the aqueous and ethanol phases were equilibrated to room temperature prior to use. The mRNA-loaded LNPs were generated using a microfluidic device (NanoGenerator Flex, Precigenome LLC, CA, USA) at a total flow rate of 4 mL/min, with an aqueous-to-organic phase flow rate ratio of 3:1. The initial mixture (∼0.25 mL from the first 5 seconds) was discarded to avoid the affect of non-equilibrium flow and residual buffer in the outlet channel. The primary mRNA-LNP suspension, containing 8 mM lipid and an N:P ratio of 6:1, was immediately diluted tenfold in Tris buffer (pH 7.4) to reduce ethanol concentration and facilitate buffer exchange. Ultrafiltration at 3,000 rpm using a 100 kDa MWCO Millipore filter was employed to remove excess buffer. The buffer replacement process was repeated to ensure the final ethanol content was reduced to less than 0.5%. For subsequent testing, mRNA-LNPs at various concentrations were concentrated using ultrafiltration. As a negative control, drug-free LNPs were prepared by using 100 mM sodium citrate buffer (pH 4.0) as the aqueous phase, following the same procedure. A 30 kDa MWCO Millipore filter was used in the cases of drug-free LNPs.

### Dynamic light scattering (DLS)

The hydrodynamic radius (R_h_) and polydispersity index (PdI) of drug-free LNPs and raw mRNA-LNPs were measured using a NanoBrook Omni multi-angle particle size and zeta potential analyzer (Brookhaven Instruments, NH, USA). Prior to measurement, 25 μL of the LNP samples was diluted with 500 μL of DEPC water and placed in a 50 μL disposable plastic cuvette (Brookhaven). Measurements were conducted at a scattering angle of 90° using a 640 nm laser and with the dust filter enabled. The refractive index of the particles was set to 1.45, following Ref. (12). Samples were equilibrated at 25 °C for 120 seconds before measurement. The apparent hydrodynamic diameter was determined using the Einstein-Stokes equation and is reported as the radius of an equivalent spherical particle.

### Cryogenic electron microscopy (cryo-EM)

The morphology of drug-free LNPs and raw mRNA-LNPs was examined using cryogenic electron microscopy (cryo-EM). Sample preparation was conducted with a Leica EM GP2 automatic plunge freezer (Wetzlar, Germany). Briefly, 3 μL of the LNP suspension, at a lipid concentration of 5-10 mg/mL, was applied to a plasma-cleaned lacey copper grid coated with a continuous carbon film. Excess sample was blotted without damaging the carbon layer. The grids were then stored in liquid nitrogen prior to imaging. Cryo-EM imaging was performed on a Talos F200C G2 Microscope (Thermo Fisher Scientific, MA, USA) operated at 200 kV, utilizing a Schottky thermal field emission super-bright electron gun. During imaging, sample grids were maintained below -170 °C. Images were captured using a FEI Ceta 4k × 4k camera, with acquisition facilitated by EPU and Serial EM software.

### Encapsulation efficiency

The encapsulation efficiency (EE%) of our mRNA-LNP sample was determined using the Quant-iT RiboGreen RNA Quantification Kit, following the manufacturer’s protocol, in a black 96-well plate. In brief, 1 μL of mRNA-LNP suspension was diluted in 100 μL of Tris-EDTA (TE) buffer for free mRNA measurement or TE buffer containing 2% Triton X-100 for total mRNA measurement. Encapsulated mRNA was not accessible to RiboGreen, whereas Triton X-100 disrupted the LNPs, releasing the mRNA. Subsequently, 100 μL of a 2,000-fold or 200-fold diluted RiboGreen solution was added to the respective wells containing the free or total mRNA solutions. RiboGreen fluorescence, which intensifies upon binding to nucleic acids, was measured at 525 nm (excitation at 425 nm) using a Spark multimode microplate reader (Tecan, Männedorf, Switzerland). Calibration curves, ranging from 0 to 100 ng/mL and 0 to 2,000 ng/mL of mRNA, were prepared by serial dilutions and mixed with the corresponding RiboGreen solutions in the same plate. Free and total mRNA concentrations were calculated from the calibration curves, and encapsulation efficiency (EE%) was calculated using the formula:

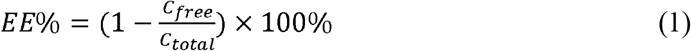

where *C*_total_ represents the total mRNA concentration in the presence of 2% Triton X-100, and *C*_free_ represents the free (unencapsulated) mRNA concentration.

### *In vitro* activity

HEK293T cells (Human Embryonic Kidney 293 Cells transformed with sheared Simian Virus 40 Large T Antigen) were obtained from the Cell Resource Center of the Chinese Academy of Sciences (Shanghai, China) and cultured following the manufacturer’s protocols. Cells were maintained in high-glucose Dulbecco’s Modified Eagle Medium (DMEM, Thermo Fisher Scientific, CA, USA) supplemented with 10% fetal bovine serum (FBS, Gibco, Australia) and 1% penicillin/streptomycin, and incubated at 37 °C in a 5% CO_2_ atmosphere. Third-passage cells, at a density of 3×10^5^ cells/mL, were seeded into 24-well plates (500 μL per well) and allowed to adhere for 24 hours prior to transfection. For transfection, 20 μL of mRNA-LNP suspension was added dropwise to each well in triplicate, followed by a 24-hour incubation. The expression of mRNA in HEK293T cells was assessed by measuring firefly luciferase activity using the Firefly Luciferase Reporter Gene Assay Kit (Beyotime Biotechnology, Shanghai, China) as per the manufacturer’s instructions. After removing the culture medium, 100 μL of cell lysis buffer was added to each well, and the cell lysates were centrifuged at 10,000×g for 5 minutes. A 40 μL aliquot of the supernatant was mixed with 100 μL of Firefly Luciferase Assay Reagent, and luminescence (RLU) was measured using a SpectraMax iD5 multi-mode microplate reader (Becton, Dickinson and Company, NJ, USA). The RLU values were normalized to the amount of encapsulated mRNA, quantified by the Quant-iT RiboGreen RNA Quantification Kit, and reported as normalized RLU (Norm RLU).

### Nano flow cytometry (NanoFCM)

The payload and capacity of mRNA-LNPs were quantified using a Flow NanoAnalyzer (NanoFCM Inc., Fujian, China). The mRNA-LNP suspension was serially diluted to achieve particle concentrations between 1×10^8^ to 1×10^10^ particles/mL in buffer solutions. These dilutions were manually processed and analyzed by the Flow NanoAnalyzer. SYTO 13, a green fluorescent nucleic acid stain (Invitrogen, Thermo Fisher Scientific, MA, USA), was added to the diluted samples and incubated at 37 °C for 15 minutes prior to analysis. Subsequently, 100 μL of the stained sample was loaded into the analyzer to assess particle concentration, the proportion of drug-free LNPs, and the payload of mRNA within the LNPs.

### Small angle X-ray scattering (SAXS)

SAXS measurements were conducted at the BL19U2 beamline of the Shanghai Synchrotron Radiation Facility (SSRF) using a synchrotron X-ray source with a wavelength of 0.103 nm. The experimental setup included a Pilatus 2M detector (172 μm×172 μm pixel size), offering a resolution of 1043×981 pixels. The sample-to-detector distance was set to 2.6 m, allowing for a scattering vector range (*q*) from 0.01 Å^-1^ to 0.43 Å^-1^. Twenty consecutive two-dimensional (2D) SAXS images were recorded with an exposure time of 0.5 seconds per frame. The resulting images were integrated into one-dimensional (1D) intensity curves using BioXTAS RAW 2.3.0 software (61). The intensity curves were averaged across all 20 frames, and solvent background subtraction was performed to obtain the final data.

### Small angle neutron scattering (SANS)

SANS experiments were conducted using both the Small Angle Neutron Scattering (SANS) instrument BL-01 (62) and the Very Small Angle Neutron Scattering (VSANS) instrument BL-14 (63) at the China Spallation Neutron Source (CSNS). The SANS data were collected in a *q* range of 0.005 to 0.6 Å^-1^, with a sample-to-detector distance of 4 m and neutron wavelengths ranging from 1 Å to 12 Å. The VSANS data were collected in a *q* range of 0.003 to 0.2 Å^-1^, neutron wavelengths ranging from 6 Å to 10.5 Å. LNP samples, diluted to a final concentration of 3 mg/mL in a solvent mixture of D_2_O and H_2_O, were placed in quartz cells with a 2 mm optical path length. Measurement durations varied from 0.5 to 3 hours, based on the scattering intensity. All experiments were conducted at 298 K. The SANS data presented were background-subtracted (solvent), corrected for empty cell contributions, and adjusted for transmission.

### Data analysis of SAXS

The scattering pattern of a particle *I*(*q*) is a Fourier transform of its pair distance distribution function, P(r), being related to each other by the equation (64):

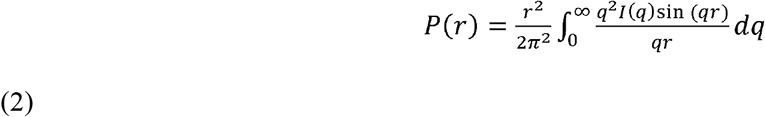

where *r* is the pair distance in real space, and *q* is the momentum transfer in reciprocal space. Furthermore, the radius of gyration (R_g_) can be calculated from the *P*(*r*) function using the equation (64):

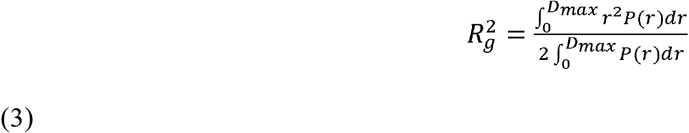

where *D*_*max*_ is the maximum dimension of the particle.

### Data analysis of SANS

The analysis of Small-Angle Neutron Scattering (SANS) and Very Small Angle Neutron Scattering (VSANS) data was performed using SasView software (https://www.sasview.org/). The lipid composition of the lipid nanoparticles (LNPs) was determined using samples with a lipid molar ratio of SM-102:DSPC:Chol:DMG-PEG2000 of 50:10:38.5:1.5, incorporating deuterated d70-DSPC and d7-Chol. The VSANS data were fitted using a core-shell ellipsoid model (41), with shell thickness constrained to be isotropic and the ratio of the equatorial to polar radius fixed at 1.2 based on cryo-EM images of drug-free LNPs and mRNA-LNPs (Figure 1b). The fitting was conducted over a *q* range of 0.003 Å^-1^ to 0.2 Å^-1^.

The following parameters were estimated and fixed:

(i) The scale factor and background were individually determined for each dataset.
(ii) The solvent scattering length density (SLD) was calculated based on the D_2_O/H_2_O ratio used, referencing the SLDs of D_2_O and H_2_O (Table S1).
(iii) Polydispersity, characterized by the Schulz distribution, was estimated from the data and fixed across all solvent contrasts.

The fitting parameters included:

(i) Constrained radii and thickness, fitted simultaneously across all solvent contrasts.
(ii) Shell and core SLDs, fitted simultaneously for each solvent contrast.

Table S2 presents the fitting parameters for the VSANS data of drug-free LNPs, and raw and adjusted mRNA-LNPs. For structural resolution of the core, the SLD of the mixed mRNA and lipid core was calculated using the formula:

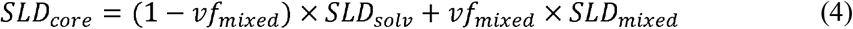

where *SLD*_*core*_ is the core SLD, *vf*_*mixed*_ is the volume fraction of mixed mRNA and lipids, *SLD*_*solv*_ is the solvent SLD, and *SLD*_*mixed*_ is the SLD of the mixed mRNA and lipids. The volume fraction of solvent (*vf*_*solv*_ ) and the dry SLD (*SLD*_*dry*_)) of the shell and the mixture of the core were derived from the fitted wet SLDs (*SLD*_*wet*_):

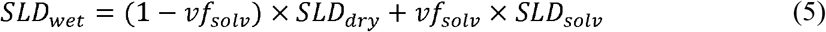

where *SLD*_*solv*_ is calculated for each contrast as detailed in Table S1. The calculated dry SLDs and solvent volume fractions for the core and shell are reported as the mean of these calculated values, with standard deviations indicating the error.

The lipid composition of core and shell was determined using the method outlined by Sebastiani et al. (25). This method partitions the molecules between shell and core based on molecular volumes and available volumes. The calculated SLDs were matched to the fitted SLDs to determine the volume fractions, which were then converted to molar fractions using component molecular volumes. The calculations assume the conservation of the input lipid molar ratio and consider the encapsulation efficiency as measured by the RiboGreen assay. All DSPC and PEG lipids were assumed to be in the shell, and all mRNA was assumed to be in the core. The conversion from SLD to volume fraction and subsequently to molar fraction is described as follows:

From SLD to volume fraction:

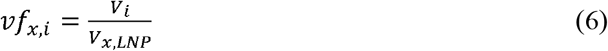

where *i* is any of the LNP components including the solvent, and *x* is either core or shell.

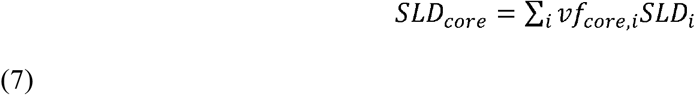

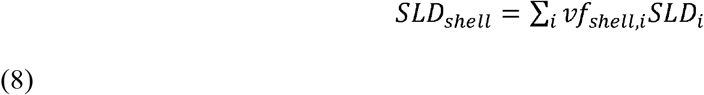

From volume fraction to molar fraction:

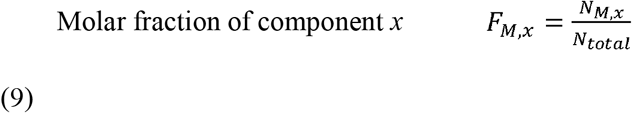

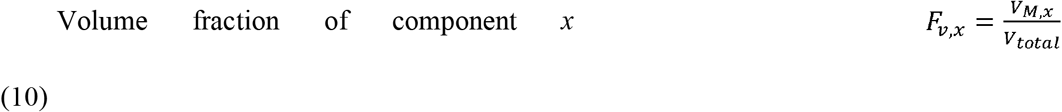

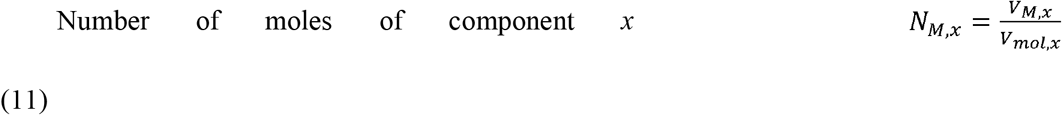

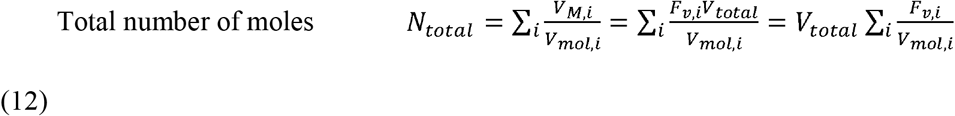

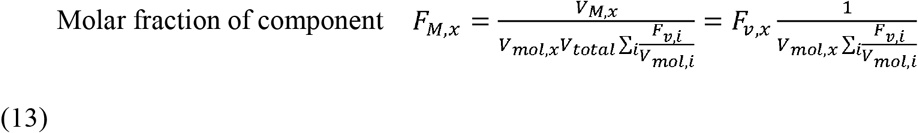

## Table of Content

Model of the mRNA-LNP sample

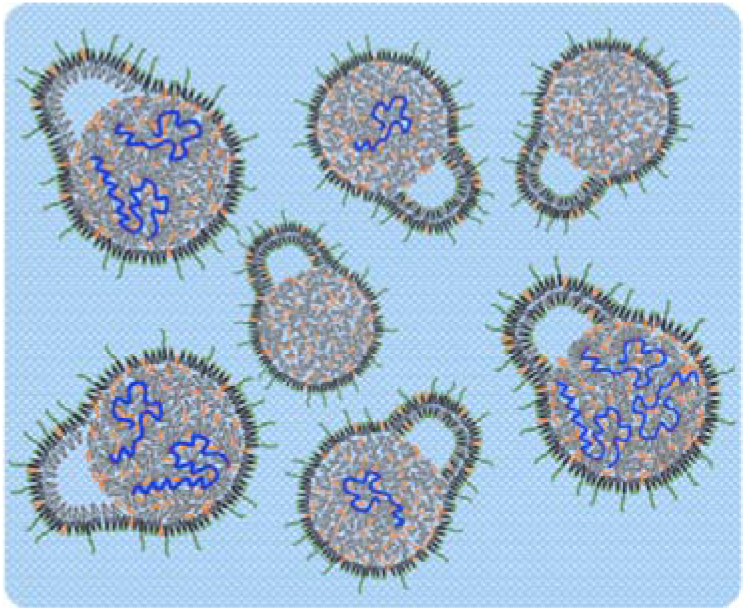

