## supporting information for "Structural Characterization of mRNA Lipid Nanoparticles in the Presence of Intrinsic Drug-free Lipid Nanoparticles"

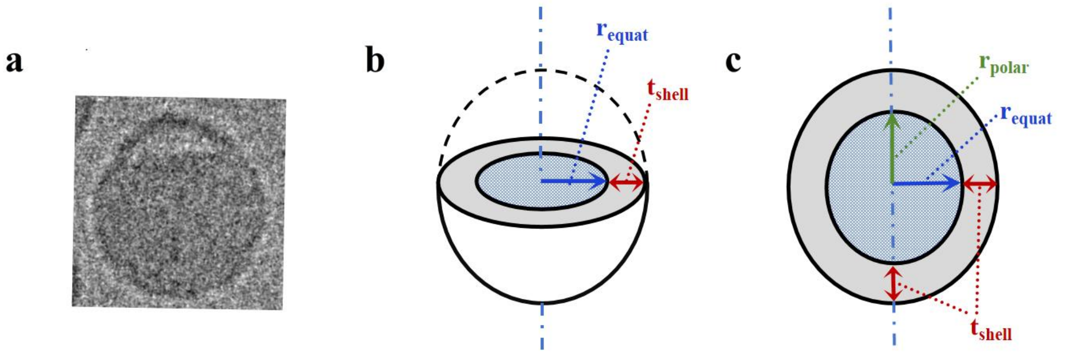


Figure S1. Core-shell ellipsoid model for fitting drug-free LNPs and mRNA-LNPs. (a) Representative cryo-EM image of mRNA-LNPs. (b) Schematic representation of a prolate ellipsoid used in the model. The diagram illustrates the geometric parameters: $r_{equat}$, $r_{polar}$, $x_{core}$, $t_{shell}$, $SLD_{core}$, and $SLD_{shell}$ are equatorial radius of core, polar radius of core, ratio between equatorial radius and polar radius, thickness of shell, scattering length density of core, and scattering length density of shell, respectively.

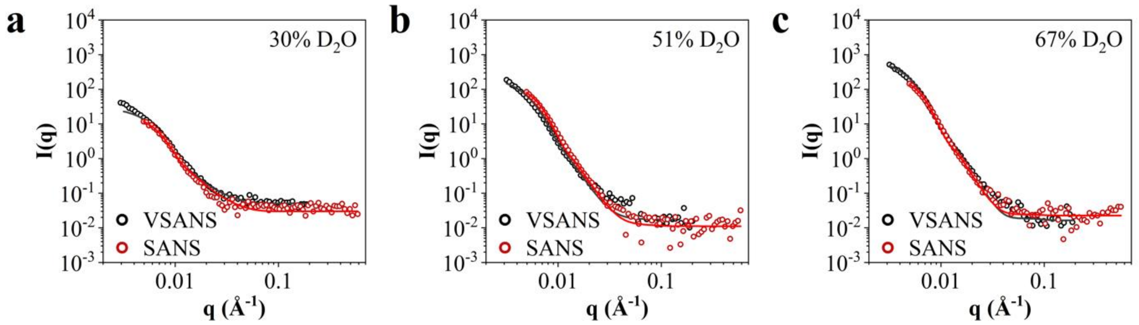


Figure S2. Comparison between VSANS (black circles) and SANS (red circles) data of drug-free LNPs in a *q* range of 0.003 Å$^{-1}$ to 0.6 Å$^{-1}$ in three different solvent contrasts: (a) 30%, (b) 51%, and (c) 67%.The fitting curves derived from our structural model of drug-free LNPs for the VSANS and SANS data are presented in black and red solid lines, respectively.

Table S1. Calculated SLDs and molecular volumes for mRNA-LNP components.

| Components | Input Formula | Molar Volume ($\times 10^{24}$ Å$^3$/mol) | Neutron SLD ($\times 10^{-6}$ Å$^{-2}$) |
|---|---|---|---|
| Water | $H_2O$ | 18.015 | -0.561 |
| Heavy water | $D_2O$ | 18.015 | 6.390 |
| SM-102 | $C_{44}H_{87}NO_5$ | 767.75 | 0.043 |
| d7-Chol | $C_{27}H_{39}D_7O$ | 364.76 | 1.416 |
| d70-DSPC | $C_{44}H_{18}D_{70}NO_8P$ | 795.56 | 5.698 |
| DMG-PEG2000 | $C_{122}H_{242}O_{50}$ | 2091.0 | 0.565 |
| DMG | $C_{32}H_{62}O_5$ | 585.35 | 0.101 |
| mRNA | $C_{27217}H_{33903}N_{11187}O_{19696}P_{2856}$ | 510801 | 3.463 |

The neutron scattering length density values (SLDs) for the water and lipid components were calculated using the NIST Neutron activation and scattering calculator (https://www.ncnr.nist.gov/resources/activation/), whereas the SLDs for the nucleic acids were calculated using the STFC ISIS Protein Scattering Length Density Calculator (http://psldc.isis.rl.ac.uk/Psldc/licence.html). Densities of above materials were provided by suppliers. In all cases, the SLD was calculated assuming 100% $H_2O$ in the solution and 0% deuteration.

Table S2. SLD value resulting from the fit of VSANS curves of drug-free LNPs, adjusted mRNA-LNPs, and raw mRNA-LNPs.

| Sample | $D_2O$ | $SLD_{core}$ ($\times 10^{-6}$ Å$^{-2}$) | $SLD_{shell}$ ($\times 10^{-6}$ Å$^{-2}$) |
|---|---|---|---|
| drug-free LNPs | 30% | 0.57 | 2.12 |
| | 51% | 0.75 | 2.67 |
| | 67% | 0.95 | 3.27 |
| | 100% | 1.44 | 4.37 |
| adjusted mRNA-LNPs | 30% | 0.78 | 2.15 |
| | 51% | 1.25 | 2.71 |
| | 67% | 1.53 | 3.38 |
| | 100% | 2.33 | 4.52 |
| raw mRNA-LNPs | 30% | 0.69 | 2.14 |
| | 51% | 1.04 | 2.70 |
| | 67% | 1.36 | 3.28 |
| | 100% | 1.92 | 4.46 |

Note that raw mRNA-LNPs refer to the experimental mRNA-LNP samples, comprising both drug-free LNPs and mRNA-loaded LNPs. In contrast, adjusted mRNA-LNPs represent mRNA-LNPs isolated from these samples by excluding the contribution of drug-free LNPs, containing only LNPs loaded with mRNA.
